# Cell-Type Specific Single-Cell Signatures Reveal Nephrotoxic Drug Affects

**DOI:** 10.1101/2025.06.17.660070

**Authors:** Aditi Kuchi, Jose Miguel Acitores Cortina, Hongyu Liu, Yasaman Fatapour, Jacob Berkowitz, Nicholas P. Tatonetti

## Abstract

Drug-induced acute kidney injury (AKI) affects about 20% of hospitalized AKI patients, a significant contributor to morbidity and mortality. The lack of understanding of the kidney system and functioning of nephrotoxic drugs contributes to hospital-acquired AKI cases. AKI is difficult to predict because of its complex injury mechanism and the numerous pathways through which it manifests. Traditional toxicity biomarkers, like elevated creatinine levels, detect AKI only after significant kidney injury has occurred. Concurrently, advancements in single cell RNA sequencing (scRNAseq) have improved our ability to map cellular heterogeneity within tissues, potentially enabling the study of drug effects at a single cell level. We hypothesized that only particular subtypes of kidney cells may be responsible for observed nephrotoxicity and explain prediction challenges. To test this, we generated cellular response scores for 32 kidney cell types from the Human Cell Atlas and estimated drug effects. We identified significant expression differences in 6 cell types (e.g. Indistinct intercalated cell p = 0.009, Epithelial Progenitor cell, p = 0.04). We also developed an ensemble model that achieved an AUROC of 0.6 across different kidney cell populations - a significant improvement over using traditional bulk RNA sequencing alone. The single-cell transcriptomic signatures we identified potentially reveal unexplained molecular mechanisms of nephrotoxicity.

**Author Summary:** The prediction and early detection of drug-induced kidney injury is a significant clinical challenge since physicians rely on biomarkers that only become elevated after substantial kidney damage has occurred, limiting opportunities for intervention and patient protection. Our investigation utilized single-cell data and available drug toxicity information to examine how individual kidney cell populations respond to potentially harmful medications. We hypothesized that specific kidney cell subtypes are primarily responsible for observed drug toxicity, which may explain the difficulties in predicting drug-induced kidney injury.

Through comprehensive analysis of 32 distinct kidney cell types, we identified six specific cellular populations that demonstrate differential responses to nephrotoxic compounds. We subsequently developed models that demonstrate superior predictive performance compared to analytical approaches using bulk RNA sequencing data. Our methodology represents a substantial advancement in precision medicine approaches to drug safety. These findings have important implications for clinical practice and patient safety. The cellular signatures we identified may enable earlier detection of kidney injury risk, potentially allowing clinicians to modify treatment regimens before irreversible damage occurs. Our work establishes a foundation for improved drug safety protocols and may contribute to reducing medication-related kidney injury in hospitalized patients.

## 1. Introduction

Acute Kidney Injury (AKI) is a critical health concern, accounting for 20% to 60% of AKI cases in hospitalized patients ^1^. The incidence of AKI in hospitalized patients ranges from 5% to 67% in patients admitted to intensive care ^2^. A significant contributor to AKI is drug-induced nephrotoxicity, which is particularly concerning due to the kidney’s crucial role in excreting toxic metabolites and drugs. This risk is further elevated in older age groups who often have coexisting morbidities and are exposed to more diagnostic procedures and therapies^3^.

Drug-induced nephrotoxicity operates through complex mechanisms that affect specific cellular components of the kidney. As detailed by Shi et.al., ^4^ nephrotoxic agents can be categorized based on the histological component affected, including proximal tubular injury (PTI), tubular obstruction by crystals or drug metabolites, and interstitial nephritis. At the cellular level, drug toxicity manifests through several distinct pathways that damage kidney cells, such as accumulation of nephrotoxic agents in proximal convoluted tubule epithelial cells, causing cellular damage, disruption of ATP production, triggering cell death, induction of oxidative stress and activation of ferroptosis^5^ (iron overload) pathways in acute kidney injury cases. These mechanisms represent key cellular processes that contribute to nephrotoxicity across various drug classes.

Current bulk RNAseq studies have mostly focused on animal models to find toxicity biomarkers^6^. Despite major consortiums like the ILSI/HESI Nephrotoxicity working group identifying promising panels of biomarkers through collaborative research, translational challenges persist between clinical outcomes and pre-clinical predictions. While these studies have developed expression biomarkers in rodent models, there remains a significant gap in translating these findings to human patients, with only limited assays being employed by pharmaceutical companies to assess nephrotoxicity before deploying drugs to market. The high levels of discrepancy between the low failure rate of drug candidates due to nephrotoxicity in pre-clinical trials (only 7%)^7^, and the high incidence of AKI in intensive care patients (20-50%)^8^ brings to light the critical need for more detailed and informed analysis on toxicity data.

Recent advancements in single-cell RNA sequencing (scRNAseq) technology have advanced our understanding of cellular heterogeneity within the human kidney, providing unprecedented resolution in studying drug effects at the cellular level. The abundance of human single-cell data in kidney research has grown exponentially in recent years. For instance, the Human Cell Atlas project ^9^ has analyzed over 100,000 cells from human kidney tissue, providing a high-resolution map of human kidney cell types and states which we use in this study. This abundance of human cell-level data, combined with comprehensive information on drug targets, presents an opportunity to identify if particular subtypes of kidney cells are responsible for nephrotoxic effects of drugs.

This hypothesis is not without precedent. Recently, Muto et al. (2021) ^10^ used single-cell transcriptomics of human kidney organoids to identify specific proximal tubule cell populations that are particularly vulnerable to cisplatin-induced injury ^11^, a common consequence of chemotherapy. Similar analyses, when extended to other drug-induced injuries, could reveal cell type-specific vulnerabilities and resistance mechanisms in human kidney tissue. The integration of drug target information with single-cell data also opens up new avenues for predictive drug toxicity in patients.

In this work, we analyze kidney specific bulk RNAseq data from LINCS ^12,13^, and single-cell RNAseq data from Human Cell Atlas (HCA)^14^ in combination with curated datasets with known drug toxicity annotations derived from labels (Ryan reference set^15^, ACS FDA labels^16^, DIRIL dataset^17^) to show that single cell data 1) has more statistical power than bulk RNAseq dataset to create a toxicity signature for drugs, 2) certain cell types that could be highly affected by nephrotoxic drugs are masked in bulk RNAseq representation, and 3) the presence of such a cell-type specific toxicity signature could help isolate drug candidates whose mechanisms are not yet known to cause nephrotoxicity. Our findings highlight the presence of a cell type that is most susceptible to drug induced injury, consistent with other literature in the field^18^.

## 2. Results

### 2.1 Simulations show bulk RNA data has limited ability to identify nephrotoxicity mediated by cell subtypes

In this section, we explored the use of bulk RNA sequencing for detecting nephrotoxicity signatures under the assumption that the nephrotoxicity is mediated by a kidney cell subtype. To evaluate the data in this context, we simulated scenarios where nephrotoxicity was mediated by Nephron, Endothelium, Immune and Stroma subtypes, representing approximately 82.4%, 7.8%, 7.1% and 2.6% of kidney cells, respectively. We then created pseudo-bulk data by computing the average across the cellular subtypes (see Methods). We evaluated the ability of this pseudo-bulk RNA sequencing data to differentiate nephrotoxic from non-nephrotoxic effects assuming a hypothetical nephrotoxic biomarker expression change by simulating a 0.01 to 10-fold change between treated and untreated samples. As shown in Figure 2 and 3, we found that statistical power was under 60% even at a maximum fold change of 10X for two cell-type abstraction levels.

**Figure 1.**
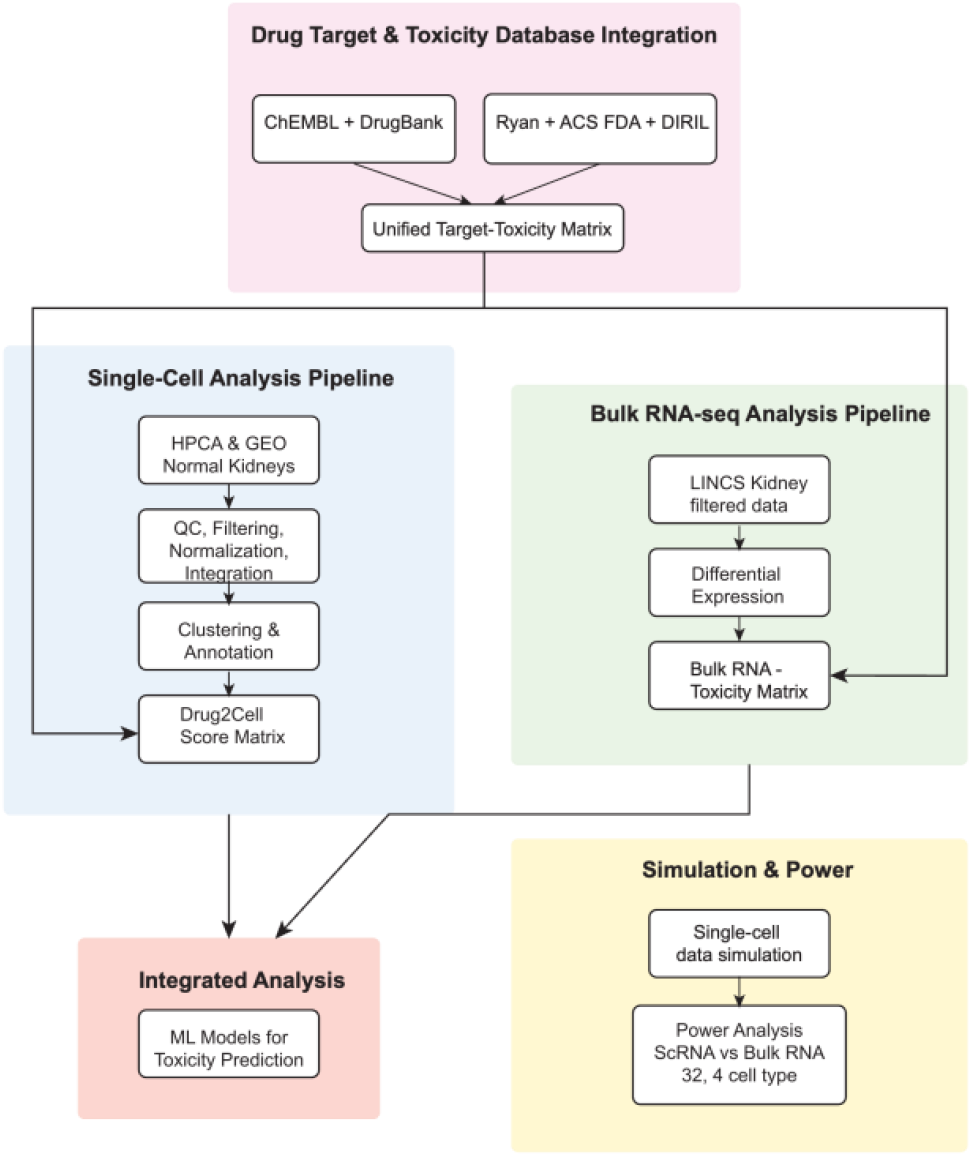
Workflow diagram – Drug target information is collected from ChEMBL^25^ and Drugbank^24^ and consolidated. Ryan reference^15^, ACS FDA label data^16^ and DIRIL datasets^17^ are combined to extract toxicity labels for drugs. Logical OR is used to label them to maximize the number of nephrotoxic drugs. To create a unified Target-Toxicity database the targets and toxicity labels are combined. This database is used in all downstream analyses. For the single-cell analysis, data is collected from HCA and GEO, selecting only healthy kidney samples. We use CellRanger to process FASTQ files, Seurat to analyze the data, QC, filter and integrate (see Methods). We annotate the clusters using SingleR, with the Kidney Cell Atlas as reference at two levels of abstraction (4 cell types, called abstract, and 32 cell types called detailed.) We analyze both levels of abstraction at all phases downstream. We then use Drug2Cell package to extract drug scores based on the target genes in the single cell matrix. We log these scores to ensure normal distribution. Our bulk RNAseq analysis extracts kidney-specific LINCS data to identify differentially expressed genes and extractable toxicity signatures. Independently, we simulated drug-affected single-cell expression data and pseudo-bulked it to compare data type power for toxicity predictions. We then used both single-cell and bulk RNAseq toxicity matrices to train machine learning methods for identifying drug toxicity signatures.

**Figure 2.**
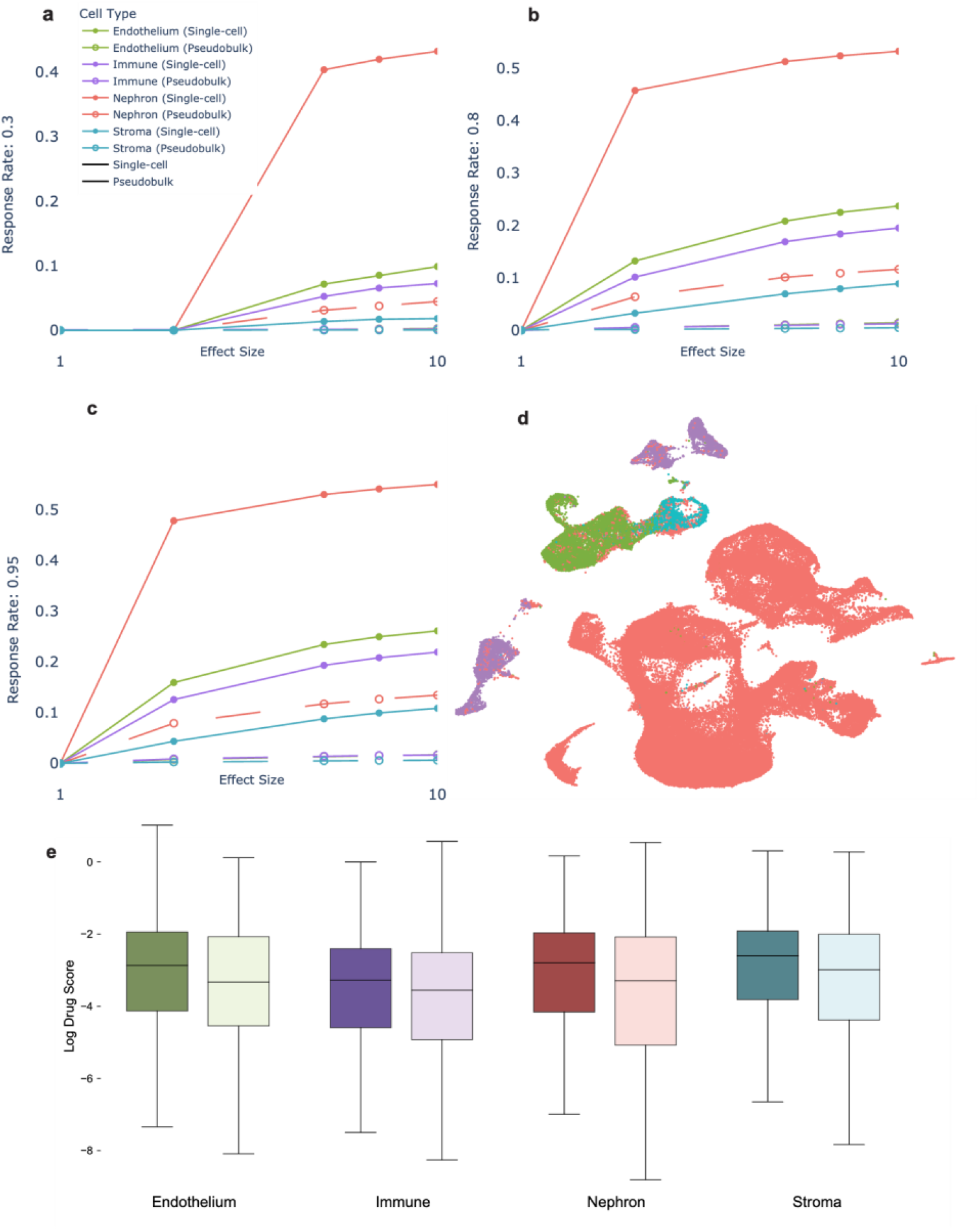
4cell-type results. In figures a), b), c) are power plots for different response rates. Response rates are defined as the percentage of cells within each sub-type that respond to a drug. Power is defined as the number of differentially expressed genes divided by total number of genes, represented on the y axis of the plots. On the x axis is the effect size. Figure d) shows a UMAP of the clustered single cell data coded by color. Many of the cells are Nephrons (82.4%). Boxplots showing the logged drug scores are in figure e). There is a marked difference in the drug scores between nephrotoxic (dark) and non-nephrotoxic (light) drugs. The nephrotoxic drug scores trend higher.

**Figure 3.**
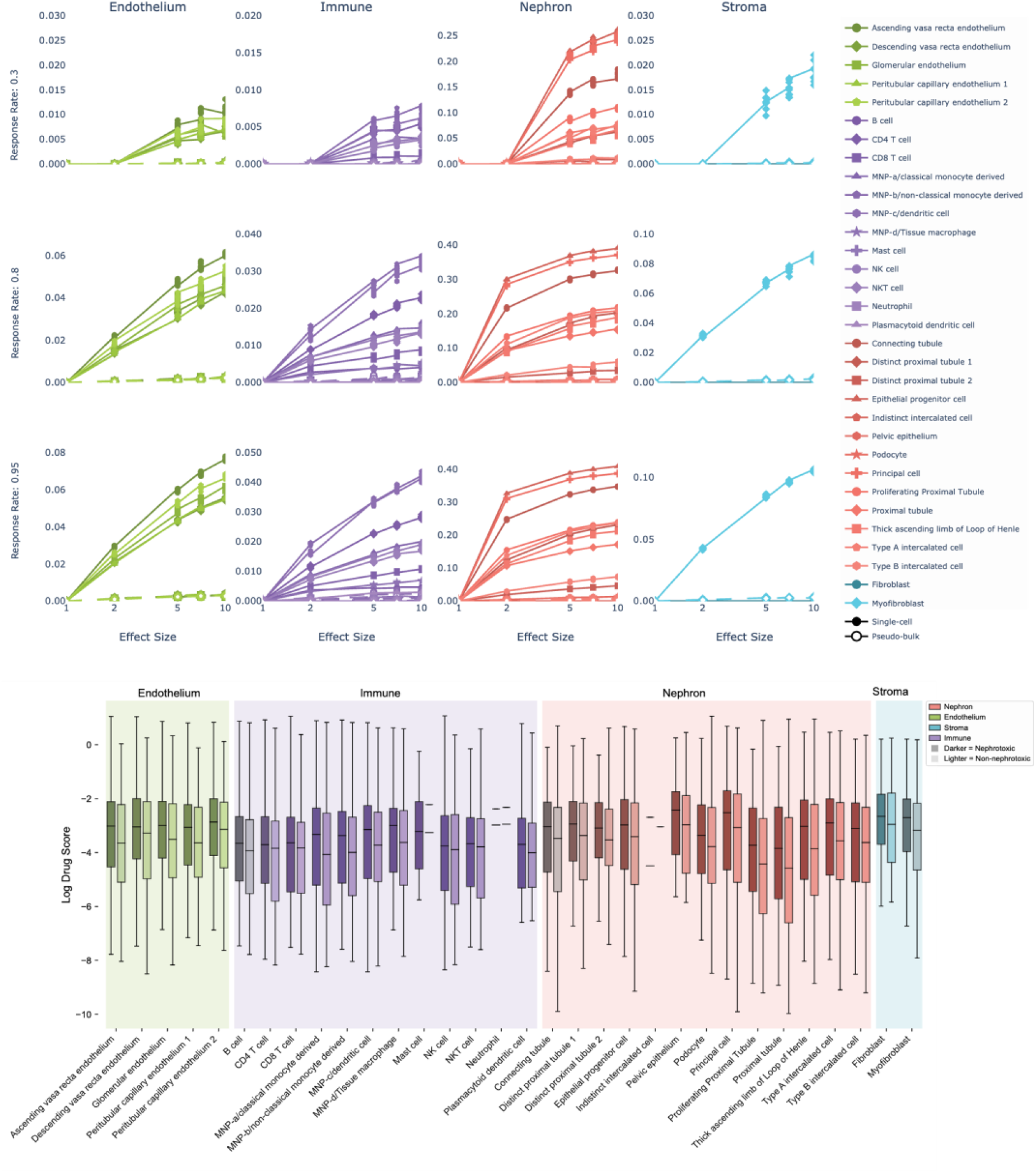
Plots showing single cell data having higher nephrotoxicity detection power compared to pseudo-bulked data. Box plots showing the increased drug scores of nephrotoxic drugs (dark) in 32 cell types.

Our analysis demonstrates a positive correlation between effect size and statistical power, as illustrated in figures 2 a, b, c and 3 a. Assuming the effect size multiplier represents a directly proportional drug action, we observe larger effect sizes in the simulated data consistently yielded increased statistical power. This supports our hypothesis that single cell level drug effect studies are superior when compared to bulk data analysis for detecting toxicity signatures.

### 2.2 Nephrotoxicity Signatures are Obscured in RNAseq of kidney epithelial cells

We performed differential expression analysis on the LINCS L1000 ^12^ dataset to compare 355 nephrotoxic samples (from 19 nephrotoxic drugs) and 16,955 non-nephrotoxic samples (from 857 non-nephrotoxic drugs). The drugs were annotated based on a curated dataset including manually verified FDA drug labels from the Ryan reference set^15^, ACS FDA labels^16^ and DIRIL^17^, identifying toxicity status for kidneys. We identified 100 differentially expressed genes (p<0.05, |log fold change| <= 1)(Top 5 shown in Table 1, all in Supplemental Table 1). Independently, we trained three machine learning classifiers and evaluated using five-fold cross validation (Table 2).

**Table 1.**
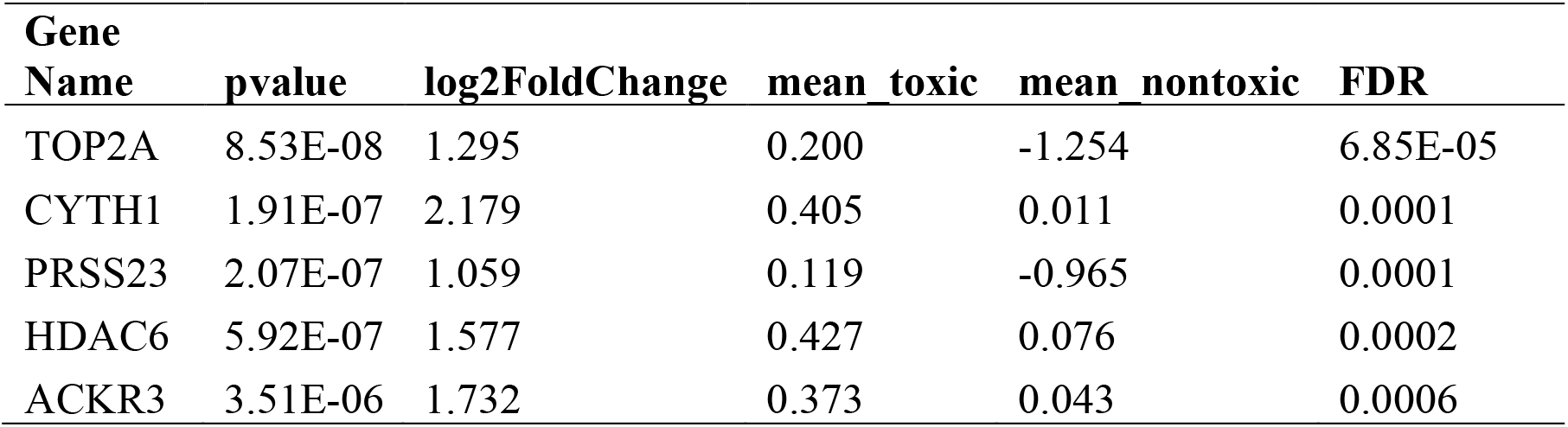
Top 5 differentially expressed genes between nephrotoxic and non-nephrotoxic drugs in kidney specific cell lines (HA1E) from LINCS. Some of these genes (HDAC6, ACKR3) are implicated in AKI and kidney damage pathways. The full list of genes can be found in Supplemental Table 1. When we collapse replicates, there are no differentially expressed genes.

| Gene Name | pvalue | log2FoldChange | mean toxic | mean nontoxic | FDR |
| --- | --- | --- | --- | --- | --- |
| TOP2A | 8.53E-08 | 1.295 | 0.200 | -1.254 | 6.85E-05 |
| CYTH1 | 1.91E-07 | 2.179 | 0.405 | 0.011 | 0.0001 |
| PRSS23 | 2.07E-07 | 1.059 | 0.119 | -0.965 | 0.0001 |
| HDAC6 | 5.92E-07 | 1.577 | 0.427 | 0.076 | 0.0002 |
| ACKR3 | 3.51E-06 | 1.732 | 0.373 | 0.043 | 0.0006 |

**Table 2.** Performance metrics for three machine learning models on kidney-specific cell lines. We used 5-fold cross validation to evaluate the models. The performance does not cross 0.5, showing the inadequacy of these models using bulk RNAseq data.

| Model | Accuracy | F1 score | Precision | Recall | AUC |
| --- | --- | --- | --- | --- | --- |
| Logistic regression | 0.458 | 0.5 | 0.4 | 0.66 | 0.416 |
| XGBoost | 0.5 | 0.0 | 0.0 | 0.0 | 0.5 |
| SVM | 0.458 | 0.5 | 0.4 | 0.66 | 0.416 |

### 2.3 Single-Cell Analysis Identifies Novel Nephrotoxic Signatures

#### 2.3.1 Nephrons are Significantly Affected by Nephrotoxic Drugs

We used single cell data from public repositories to identify novel nephrotoxic signatures. We process all datasets using CellRanger ^19^ and Seurat ^20^, following standard protocols (see Methods). We annotated the data using SingleR ^21,22^ with the kidney cell atlas^23^ as reference. We derive the drug target information from Drugbank ^24^ and ChEMBL ^25^, whereas the toxicity labels are collected from the Ryan reference, ACS FDA labels, and the DIRIL dataset. We map the cells to drugs using their targets. Using drug2cell with our custom database generates a composite score for each cell-drug pair representative of their action.

The output of the drug2cell package is a vector for each drug of the drug’s estimated effect on each cell. We grouped cells by their annotated type and then tested for differences in the drug2cell scores between scores for nephrotoxic drugs and non-nephrotoxic drugs using a T-test. We performed this evaluation at two levels of cell type abstraction (4 cell types called “abstract”, and 32 cell types called “detailed”). The 4-type abstraction is a superset of the 32-type annotations. This dual-level analysis was conducted to determine whether nephrotoxicity signals were dependent on cell type resolution. The p-values and other statistics in abstract and detailed cell type are reported in Tables 3 and 4 respectively. Supplemental Table 2 reports the p-values of all detailed cell types. We found that Nephron cells showed significant differences in drug scores between nephrotoxic and non-nephrotoxic drugs (Table 3). We found that Indistinct intercalated cell, MNP-b/non-classical monocyte derived cell, Peritubular capillary endothelium, Distinct proximal tubule, Proliferating Proximal Tubule, and Epithelial progenitor cell types showed significant differences in drug scores in the detailed cell type analysis (showed in Table 4). We found that an extra trees model trained on the detailed 32 cell types data achieved an AUROC of 0.6.

**Table 3.**
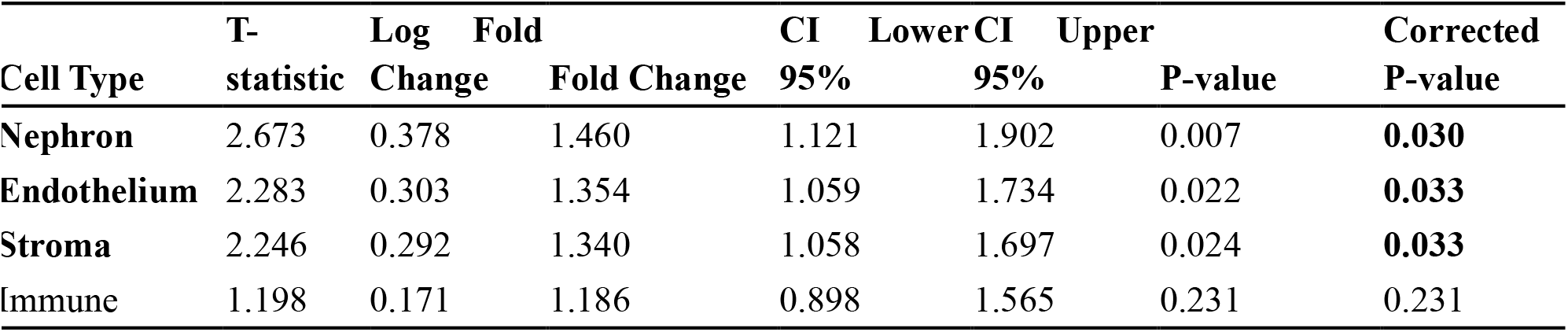
Abstract (4) cell-type t-test results show that Nephrons, Endothelium and Stroma are significant.

**Table 4.**
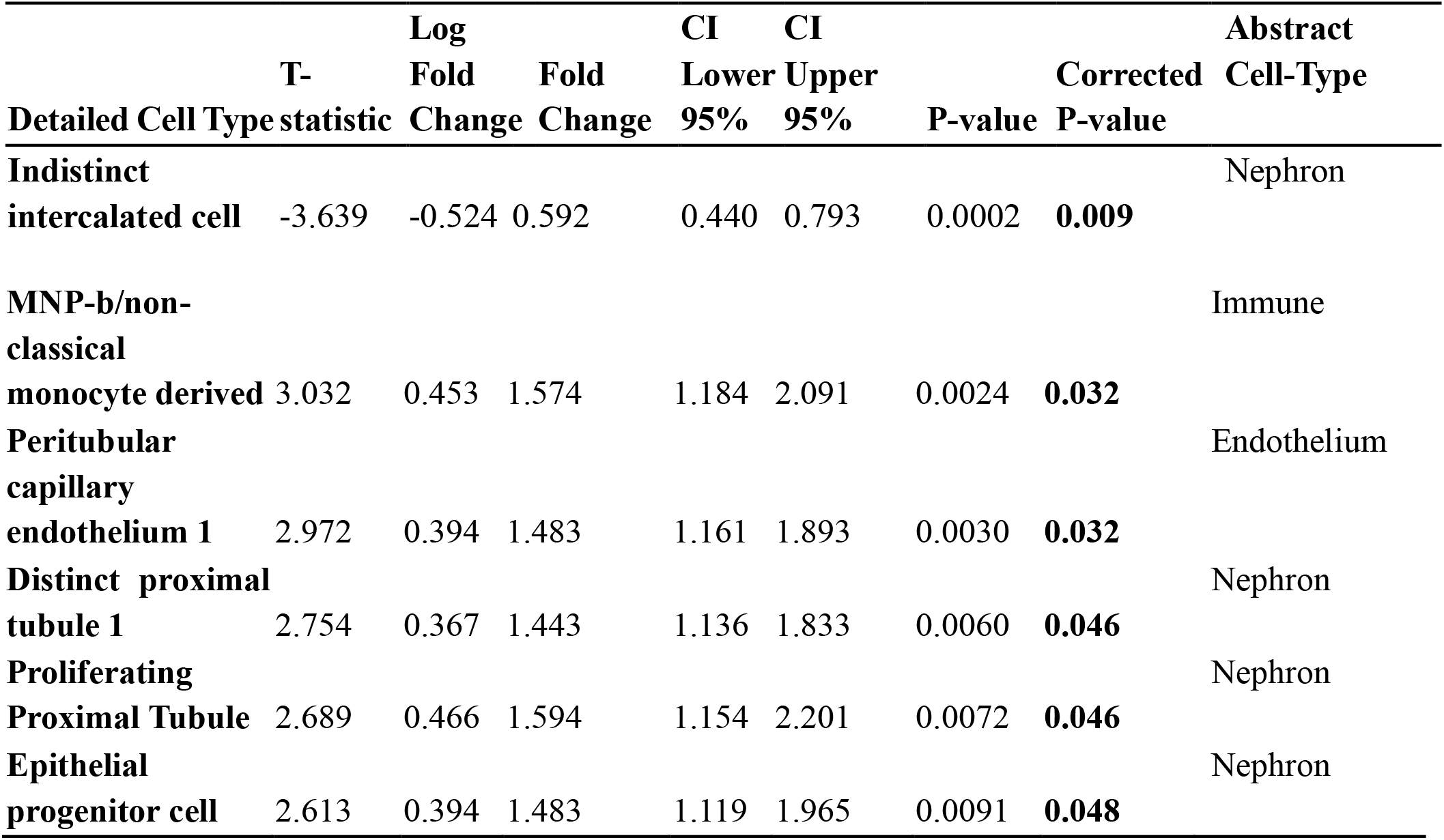
Detailed (32) cell-type t-tests shows six cells with significant expression between nephrotoxic and non-nephrotoxic drugs.

#### 2.3.2 Extra Trees Model Captures Better Toxicity Signature in Single-Cell

We evaluated 6 classical machine learning algorithms and a multilayer perceptron through ROC curve analysis, demonstrating improvement over traditional bulk RNA-seq-based models in Table 2. Individual algorithms (Logistic Regression, XGBoost, Random Forest, and Support Vector Machine, Extra Tree, Gradient Boosting) showed moderate performance with AUROC values ranging from 0.47 to 0.6 (Table 5). Single-cell data improved model performance across F1 score, sensitivity, and specificity metrics compared to bulk RNA-seq models, though AUROC values increased (bulk RNA-seq: 0.46 for Logistic Regression, SVM, 0.5 for XGBoost).

**Table 5.** Performance metrics for various machine learning models using single-cell RNAseq drug scores as features. There is a marked increase in the AUC compared to bulk RNAseq.

| Model | Accuracy | F1 Score | AUC | Sensitivity | Specificity | Lower 95% CI | Upper 95% CI |
| --- | --- | --- | --- | --- | --- | --- | --- |
| Extra Trees | 0.754 | 0.864 | **0.6** | 0.953 | 0.162 | 0.495 | 0.705 |
| Gradient Boosting | 0.731 | 0.864 | 0.597 | 0.921 | 0.162 | 0.488 | 0.693 |
| Ensemble | 0.760 | 0.869 | 0.594 | 0.953 | 0.186 | 0.483 | 0.691 |
| Random Forest | 0.771 | 0.869 | 0.591 | 0.968 | 0.186 | 0.480 | 0.698 |
| XGBoost | 0.731 | 0.869 | 0.564 | 0.882 | 0.279 | 0.450 | 0.674 |
| SVM | 0.748 | 0.856 | 0.554 | 1 | 0 | 0.458 | 0.660 |
| Logistic Regression | 0.725 | 0.856 | 0.479 | 0.968 | 0 | 0.373 | 0.584 |
| Multilayer Perceptron | 0.432 | 0.856 | 0.496 | 1 | 0 | 0.389 | 0.599 |

## 3 Discussion

Our study used single-cell data to investigate the association between specific kidney cell types and nephrotoxic drugs. The results revealed statistically significant associations between nephrons, endothelium and immune cells with nephrotoxic drugs. Specifically, we identified six detailed cell types that demonstrated significance: Indistinct intercalated cell, MNP-b/non-classical monocyte derived cell, Peritubular capillary endothelium, Distinct proximal tubule, Proliferating Proximal Tubule, and Epithelial progenitor cells. Notably, the majority of these are nephrons, suggesting that nephron-specific cellular mechanisms play a crucial role in drug-induced kidney injury.

The abundance of single-cell data and the inherent heterogeneity of kidney tissue allow for a detailed examination of specific regions that may be more susceptible to drug effects. The kidney is an ideal organ for this analysis due to several important characteristics. It contains discrete cell types that perform distinct functions in drug metabolism and excretion. Different anatomical regions of the kidney are specifically responsible for metabolizing various drugs, making it possible to pinpoint where toxicity may occur. A granular understanding could impact the drug screening process. Since kidneys are responsible for filtration and excretion of drug metabolites, the byproducts are easily measurable in urine and blood, providing convenient biomarkers for toxicity assessment. This approach is also readily extensible to other tissue types and diseases, with liver tissue being another excellent candidate for similar analyses due to its central role in drug metabolism. If we can identify a stronger signature indicative of toxicity in certain cell types, it could lead to the development of more accurate models for screening drug candidates. Moreover, this knowledge could benefit patients undergoing intensive care who are at risk of further complications.

Our analysis reveals critical insights into drug-induced kidney injury that are obscured in bulk RNA sequencing studies. While bulk transcriptomics captures general tissue responses, it masks cell type-specific signals due to averaging across the kidney’s cellular heterogeneity. This “signal dilution” limits detection of toxicity patterns confined to particular microenvironments, or what we term “toxicity niches.” In contrast, single-cell RNA sequencing preserves this cellular resolution. Even with a coarse classification into four major cell types, our model demonstrates superior predictive power relative to the bulk LINCS dataset—indicating a qualitative shift in our ability to detect nephrotoxicity. This advantage likely reflects the biology of injury itself: nephrotoxic compounds often target cellular processes enriched in specific cell types. For example, proximal tubule cells are vulnerable to mitochondrial toxins due to their high metabolic activity, while podocytes are more susceptible to compounds disrupting cytoskeletal integrity. By resolving these cell type-specific responses, single-cell analysis captures toxicity signatures that bulk approaches miss. This aligns with recent studies showing distinct injury responses across kidney cell types ^18^ and underscores the importance of selective cell death and fibrosis pathways in the progression from acute kidney injury (AKI) to chronic kidney disease (CKD) ^26,27^. Ultimately, maintaining cellular context may offer valuable sensitivity in detecting early or subtle nephrotoxic events, where conventional biomarkers fall short.

One advantage of our approach is that it bypasses the need for complex pathway analysis while still capturing the essential biology of nephrotoxicity. Rather than requiring detailed knowledge of specific molecular pathways disrupted by each potential toxic drug, which varies widely between drug classes, our method leverages the cellular expression patterns and their interaction with the drug targets themselves as an integrated score of toxicity. This represents a robust approach that can detect novel forms of nephrotoxicity even when the precise mechanism is unknown. In the drug toxicity testing pipeline, focusing on these significant cell types will help pharma companies develop more targeted and precise screening methods. The power analysis we conducted specifically shows the value of utilizing single cell data to reveal nephrotoxicity signatures. Despite the promising results, our study has several limitations. There is scope to improve the simulation of single-cell data, as we do not account for functional relationships between genes. Gene expression of certain significant genes likely varies in combination with others, creating complex patterns that our current models may not fully capture. Another key limitation was the exclusive use of healthy donor kidneys. This restricts our understanding of how drug effects might manifest in diseased states. Although a study based solely on normal kidneys provides valuable baseline information, incorporating data from diseased kidneys would prove highly beneficial, making it more directly clinically relevant.

This work lays the foundation for a more nuanced and effective approach to predicting and mitigating adverse drug effects in the kidney. By continuing to refine and validate these methods, we aim to develop tools that can significantly improve drug safety assessment and ultimately enhance patient outcomes through reduced incidence of drug-induced kidney injury.

## 4 Data and Methods

### 4.1 Data Sources and Workflow

We collected and processed normal kidney single-cell RNA sequencing (scRNA-seq) data from the Human Cell Atlas (HCA) and Gene Expression Omnibus (GEO), accession number GSE131685^28^. Following the workflow shown in Figure 1, we processed and integrated these datasets using Seurat v5.1.2 in R v4.4.2. For samples with available FASTQ files, we used Cell Ranger v1.1.0 (provided by10X Genomics) to generate gene-barcode matrices. We then converted the generated gene-barcode matrices to count data and then Seurat objects.

We implemented quality control following established best practices for single-cell RNA sequencing data analysis. We filtered cells using the following criteria: (1) minimum UMI count threshold of 500; (2) minimum gene detection threshold of 250 genes per cell; and (3) log10 genes per UMI ratio greater than 0.85, which helps identify and remove potential technical artifacts. We then filtered genes by retaining only those present in at least 10 cells to ensure reliable downstream analysis. Notably, we did not implement specific doublet detection algorithms or upper thresholds for UMI or gene counts. This decision was based on methodological considerations regarding doublet detection tools, which can inadvertently remove cells with intermediate or transitional phenotypes while being optimized for datasets with discrete cell types. Instead, we opted to evaluate potential doublets during later stages of analysis by examining whether marker genes from multiple distinct cell types co-occurred in specific clusters.

We normalized the filtered data using the LogNormalize method. We identified highly variable genes using the variance-stabilizing transformation (VST) approach, selecting the top 2,000 genes for downstream dimensional reduction and clustering. Then, we applied scaling to these genes.

We first merged the data and then performed the integration step based on common cell types and genes between all samples, merging data from 24 samples and creating a comprehensive kidney cell dataset. We followed the standard Seurat integration workflow, but using reciprocal principal component analysis (RPCA) to identify common sources of integration between the datasets. We then performed principal component analysis (PCA) on the integrated data for dimensionality reduction. We determined the optimal number of principal components for downstream analysis using elbow plots. We then used these principal components as input for Uniform Manifold Approximation and Projection (UMAP) to generate two-dimensional visualizations of the cell landscape. We performed clustering using Seurat’s FindClusters function, with resolution parameters adjusted to obtain two levels of granularity in cell type identification (4 and 32 levels of abstraction). To annotate the identified clusters, we employed SingleR for automated cell type prediction, using the Kidney Cell Atlas as a reference ^23^. This approach yielded two levels of cluster resolution: a broad classification identifying 4 major cell types (endothelial cells, immune cells, nephrons, and stroma), and a finer resolution distinguishing 32 specific cell subtypes.

### 4.2 Drug Response Simulations Validate the Necessity of Cell-Level Data

To see if there is a marked difference between a baseline dataset and drug-affected single cells, we simulate single-cell data by randomly sampling 10% of the cells from the kidney normal dataset while preserving the original cell type distributions (prevalence). Drug effects were simulated by applying varying effect sizes to the baseline gene expression values, maintaining the proportion of cell type specific response rates. Effect size, in this experiment, are values ranging from 0.001 to 10 on a log scale. These are added as a multiplier to the original data to change their values by this factor. The percentage of cells that this multiplication factor is applied to are calculated based on the cell type response rate. Cell type response rate is one of 0.8 which is the assumed standard expected rate of response of a drug, 0.95 when the response rate is much higher than usual, and 0.3 to make sure to include drugs that could have very low response rates among different cell types.

The pseudocode to calculate the altered gene expression can be found in Algorithm 1. In a randomly sampled 10% dataset from the original single cell gene expression: For cell type A, with a prevalence of 80%, we select response rate (0.8, 0.95, or 0.3) number of cells. To this proportion of cells, we multiply a 1 + effect size [0.001, 10] factor to alter the gene expression.

#### Algorithm 1

Simulation of Drug Affected Single-Cell Data

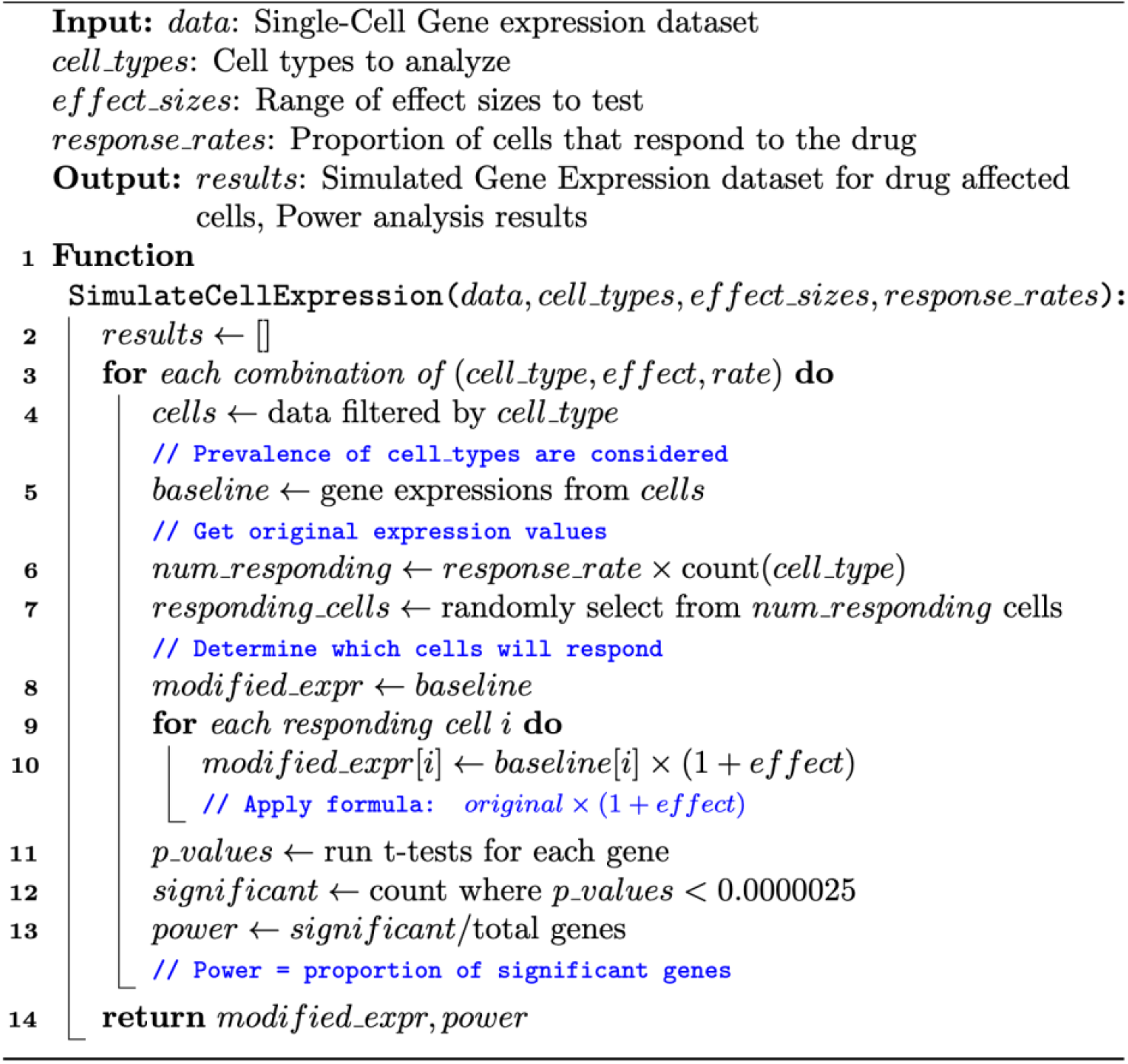

This generates a simulated single cell matrix with the same prevalence make-up as the original data, while modifying gene expression of the drug-affected cells.

To perform the power analysis on the simulated single cell data, we first pseudo-bulk it by averaging the altered gene expression values across all cell types per gene. This results in a gene expression matrix similar to what is expected in a bulk RNAseq dataset, with the cell type annotations masked. By using this data to observe the number of significant genes per effect size, and how the value increases with increasing drug effect (directly translated from the effect size value), we show that lower values of drug effects fail to provide enough statistical power to effectively exhibit patterns to measure nephrotoxicity signatures. It is also vital to note that there could be certain cell types that are highly affected by nephrotoxic drugs that are masked in bulk data, and therefore when averaged, lose their predictive power.

### 4.3 Mapping Drug Effects to Single Cells (Drug2Cell)

Once the cell type annotations from the integrated kidney single cell dataset were obtained, we furthered our analysis by employing Drug2Cell^29^, a package designed to integrate drug-target interactions with single-cell expression data as described by Kanemaru et.al. The approach combines a cell-by-gene expression matrix with a drug-by-gene target matrix to evaluate drug-target expression at the single-cell level. Drug2Cell implements a scoring system that quantifies the potential effect of drugs on individual cells based on the expression of their target genes. In our study, we applied this package with a curated dataset comprising:

A. DrugBank: We extracted information for approved or withdrawn drugs to ensure clinical relevance.
B. ChEMBL: This provided experimentally validated data for drug-target pairs.
C. Nephrotoxicity Dictionaries: We incorporated two specialized nephrotoxicity resources:
  1. Ryan reference: A manually curated list of drugs with verified nephrotoxicity status.
  2. ACS FDA labels: A comprehensive collection of FDA-approved drug labels with extracted nephrotoxicity information.
  3. DIRIL dataset: An FDA confirmed nephrotoxicity labeled dataset on 317 single-molecule drugs.

To link drug targets to their nephrotoxicity information, we combined data from DrugBank (approved drugs only) with the other three data sources. We obtain 215 nephrotoxic and 641 non-nephrotoxic drugs.

For the drug score calculation, we applied the Drug2Cell algorithm to calculate “drug scores” for each cell type identified in our integrated kidney cell dataset. We created a cell-by-gene expression matrix from our annotated single-cell data, built a drug-by-gene target matrix from our curated drug-target information, and computed the cell-specific drug scores by averaging the expression values of all genes targeted by each drug in each cell. This approach allowed us to predict and quantify potential drug responses across the various kidney cell populations, offering insights into cell type-specific drug sensitivities and potential nephrotoxicity mechanisms. To prepare our data for downstream analyses, we applied a logarithmic transformation to the generated drug scores. This transformation normalized the data distribution of the scores, reducing the impact of extreme values and enhancing data interpretability, allowing for easier comparison of drug effects across different scales. The resulting log-transformed cell-by-drug matrix was used in all downstream analyses.

### 4.4 Bulk data processing

We obtained gene expression profiles from the Library of Integrated Network-based Cellular Signatures (LINCS) L1000 dataset, which contains transcriptional responses to thousands of bioactive small molecules across multiple cell lines. For our analysis, we specifically focused on the kidney cell line HA1E to investigate kidney-specific drug responses, as detailed in our results section. This analysis was conducted independently of our single-cell approach. The raw expression data from LINCS was processed to address technical variability between replicates. We implemented a regex pattern matching approach to identify all replicates for each drug and collapsed them by averaging the expression values.

To identify gene expression signatures associated with nephrotoxicity in bulk RNA-seq data, we performed differential expression analysis comparing gene expression profiles between nephrotoxic and non-nephrotoxic drugs. As detailed in our results section, this analysis initially identified 223 differentially expressed genes when using all cell lines, which was reduced to 100 genes when restricting the analysis to kidney-specific cell lines (HA1E). However, after collapsing replicates, no statistically significant differentially expressed genes were detected, highlighting the limitations of bulk transcriptomic approaches for nephrotoxicity prediction.

### 4.5 Statistical Analysis and Machine Learning

We performed t-tests with False Discovery Rate (FDR) correction to find significant differences between nephrotoxic and non-nephrotoxic drug groups. These tests were conducted using the log-transformed drug score matrix, allowing us to identify cell types that exhibited significant differences in their responses to nephrotoxic versus non-nephrotoxic drugs. To generate the representative drug score for each cell type used in the statistical tests, we calculated the average of all drug scores pertaining to that specific cell type within our dataset.

Building upon these statistical findings, we extended the analysis by applying machine learning techniques to evaluate whether the cellular response signatures we identified could serve as effective predictors of a drug candidate’s potential nephrotoxicity.

We implemented six different algorithms: XGBoost (Extreme Gradient Boosting), Support Vector Classifier (SVC), Logistic Regression and Random Forest, Gradient Boosting and Extra Tree. In addition to these, we trained an ensemble model and an MLP.

We trained these models on our processed dataset using the log-transformed drug scores in each cell type as features and the known nephrotoxicity status as the target variable. We evaluated using 5-fold cross-validation. This approach partitions the data into k subsets, using k-1 subsets for training and the remaining subset for testing across k iterations, ensuring that all data points are used for both training and testing.

## 5 Conclusion

Our study uses single-cell RNA sequencing to identify cell type-specific signatures that elucidate mechanisms of drug-induced nephrotoxicity, particularly highlighting the vulnerability of Nephrons. These signatures promise enhanced early detection and prediction of nephrotoxicity, potentially improving drug safety screening and design. Despite limitations, like the use of only healthy donor kidneys, there is potential in expanding the dataset and incorporating diseased kidney data for validation. Future work will focus on addressing these limitations and employing advanced machine learning techniques for better predictive capabilities. This research lays a strong foundation for precise strategies in mitigating adverse drug effects on the kidney, with substantial potential to impact patient care and drug development.

## Data Availability

The original datasets used in our analysis can be accessed through their respective repositories: the Human Cell Atlas is available through DOI: 10.1038/s41586-023-05769-3), and the GSE131685 dataset can be accessed through the Gene Expression Omnibus (GEO) at https://www.ncbi.nlm.nih.gov/geo/query/acc.cgi?acc=GSE131685.

## Author Contributions

A.K. and N.P.T. conceived the study, developed the methodology, A.K. performed the analysis, created the visualizations, and wrote the original draft. N.P.T. supervised the project, provided guidance on methodology and analysis, and reviewed and edited the manuscript. J.M.A.C., H.L., Y.F., and J.B. provided advice on the research approach, reviewed and edited the manuscript. All authors read and approved the final manuscript.

## Acknowledgements

AK and NPT are supported by R35 GM131905.

